# Spatiotemporal patterns of urbanization in three Swiss urban agglomerations: insights from landscape metrics, growth modes and fractal analysis

**DOI:** 10.1101/645549

**Authors:** Martí Bosch, Jérôme Chenal

## Abstract

Urbanization is currently a global phenomenon that has become the most important form of landscape change and is increasingly affecting biodiversity and ecosystem functions. In order to evaluate the impacts of urbanization and inform urban planning, it is important to understand the spatiotemporal patterns of land use change associated to urbanization. This paper exploits three different frameworks, namely landscape metrics, urban growth modes and fractal analysis to characterize the spatiotemporal patterns of urbanization of the Swiss urban agglomerations of Zurich, Bern and Lausanne. The land use inventory provided by the Swiss Federal Statistical Office was used to assemble four temporal snapshots from 1980 to 2016 at the extent of the urban agglomerations. The time series of landscape metrics generally supports the diffusion and coalescence model of urban growth, with Zurich exhibiting most characteristics of coalescence while Bern and Lausanne seem to be at the transition between diffusion and coalescence. Nevertheless, the analysis of the urban growth modes suggest that leapfrog development occurs at all periods, which contributes to an increasing fragmentation of natural patches and maintains the fractal configuration of the landscape. The discussion reviews potential explanations for the observed landscape changes, and concludes with some planning implications.

## Introduction

The last centuries have seen an unprecedented growth of urban areas, which has resulted in dramatic conversion of natural land and profound changes in landscape patterns and the ecosystem functions that they support (Alberti 2005). The combination of current demographic prospects and the observed trends of decreasing urban densities suggest that the global amount of land occupied by cities might increase threefold (Angel et al. 2005). Quantifying urban landscape patterns in space and time is an important and necessary step to understand the driving forces and ecological impacts of urbanization (Wu 2014).

Recent decades have witnessed an increasing number of studies of the spatiotemporal patterns of land use change associated to urbanization (Dietzel et al. 2005; Seto and Fragkias 2005; Schneider and Woodcock 2008; Jenerette and Potere 2010; Wu et al. 2011; Li et al. 2013; Liu et al. 2016; Nong et al. 2018). Initial attempts to synthesis suggested that urbanization can be characterized as a two-step alternating process of diffusion and coalescence (Dietzel et al. 2005; Schneider and Woodcock 2008), nonetheless, subsequent studies challanged the empirical validity of such hypothesis. On the one hand, Jenerette and Potere (2010) examined the spatiotemporal patterns of land use change of a sample of 120 cities distributed throughout the world from 1990 to 2000, and determined that overall, urbanization leads to fragmented landscapes with more complex and heterogeneous structures. On the other hand, Li et al. (2013) determined that the two-phase diffusion and coalescence model can be over-simplistic and that urbanization might be better characterized by means of three growth modes, namely infilling, edge expansion and leapfrogging, which operate simoultaneously while alternating their relative dominance.

Paralleling the above studies, approaches from the complexity sciences have provided novel insights into the spatial organization that underpins contemporary cities (Batty 2005). Like many other complex systems, cities exhibit morphological traits that are consistent with fractal geometry and reflect the self-organizing nature of the processes that occur upon them (White and Engelen 1993). Although the scaling relationships of fractal geometry suggest the existence of strong morphological regularities over a wide variety of cities and regions (Frankhauser 1994; Batty and Longley 1994; White et al. 2015), their meaning in the context of the spatiotemporal patterns of urbanization remains unclear (Li 2000; Manson and O’Sullivan 2006; Bosch et al. 2019).

This paper intends to evaluate the spatiotemporal patterns of land use change observed in three of the largest Swiss urban agglomerations from three different perspectives, namely landscape metrics, urban growth modes and fractal analysis. The objective of this paper is twofold. The first is to assess to test whether the spatiotemporal evolution of the three agglomerations conform to the diffusion and coalescence dichotomy, the second is to explore how the three adopted perspectives complement each other. The results will serve to discuss planning implications regarding the desirability of the recent densification policies adopted in Switzerland.

## Materials and Methods

### Study area

Switzerland is a highly developed country in central Europe, with a population distributed into several interconnected mid-sized cities and a large number of small municipalities. Mainly because of the country’s topography, most urban settlements are located in its Central Plateau region, which accounts for about one third of the total Swiss territory, (42,000 km^2^) and is highly urbanized (450 inhabitants per km^2^). The Central Plateau is characterized by elevations that range from 400 to 700m, a continental temperate climate with mean annual temperatures of 9-10 xC and mean annual precipitation of 800-1400 mm, and a dominating vegetation of mixed broadleaf forest.

In line with the country’s federalist government structure, the Swiss spatial planning system is distributed between the federal state, the 26 cantons and 2495 municipalities. The federal state specifies the framework legislation and coordinates the spatial planning activities of the cantons, while cantons check the compliance of municipal development plans with cantonal and federal laws. With some exceptions, municipal administrations are in charge of their local development plans, namely the land use plan and building ordinance, and might therefore be viewed as the most important spatial planning entities. While the Federal Statue on regional planning of 1979 limited the number of new buildings constructed outside the building zones, built-up areas have since increased continuously, mainly because the municipalities can designate new building zones almost entirely autonomously (Jaeger and Schwick 2014). A major revision of the Federal Statue was accepted in 2013, which limits the amount of building zones that municipalities can designate and encourages infill development and densification by means of tax incentives. Forecasts based on the current urbanization trends predict significant increases of urban land use demands over the next decades, mostly at the expense of agricultural land located at the fringe of existing urban agglomerations (Price et al. 2015).

Given that a significant part of the cross-border urban agglomerations of Geneva and Basel (the second and third largest in Switzerland) lie beyond the Swiss boundaries (SFSO 2014), in order to ensure coherence of the land use/land cover data (see the section below), this study comprises only three of the five largest Swiss urban agglomerations, namely Zurich, Bern and Lausanne (SFSO 2018). As shown in Figure 1, the three agglomerations have undergone important population growth over the last 30 years, especially during the most recent years and at the agglomeration extent. With a total population over 1.3 million and land area of 1305 km^2^ (1038 hab/km^2^), Zurich is the largest Swiss urban agglomeration. As a leading global city and one of the world’s largest financial centers, Zurich has the country’s largest airport and railway station, and also hosts the largest Swiss universities and higher education institutions. Bern is the capital of Switzerland and fourth most populous urban agglomeration in Switzerland, with a total population of 410000 inhabitants and occupying a land area of 783 km^2^ (531 hab/km^2^). As the fifth largest Swiss urban agglomeration and the second most important student and research center after Zurich, the Lausanne agglomeration has a total population of 409000 inhabitants over a land area of 773 km^2^ (537 hab/km^2^). Given its larger population growth rate, Lausanne is likely to soon surpass Bern and become the fourth largest urban agglomeration in Switzerland. Overall, the three urban agglomerations are characterized by a pervasive public transportation system and a highly developed economy, with a 85% of the employment devoted to the tertiary sector.

**Fig. 1.**
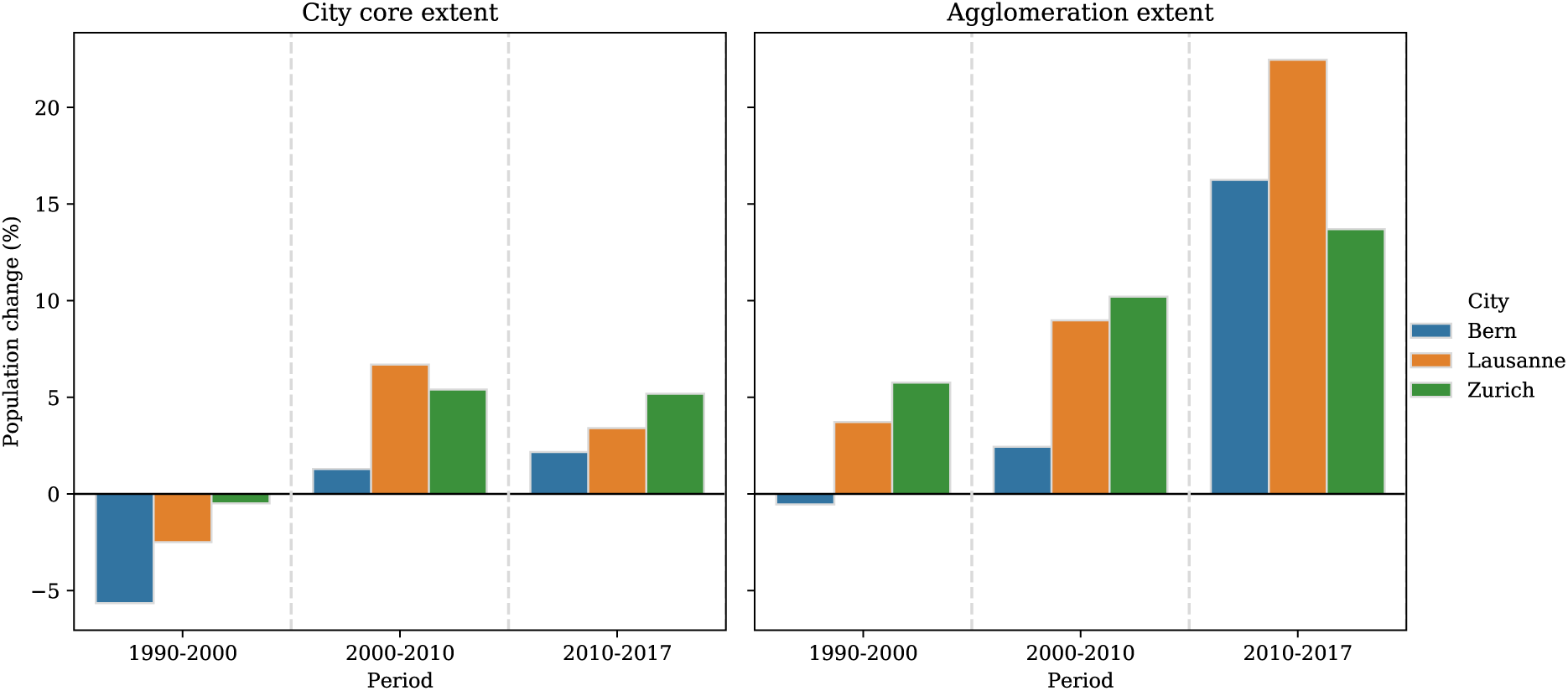
Population change of the three regions of study at the city core (left) and agglomeration extent (right) over the periods of 1990-2000, 2000-2010 and 2010-2017. Data from the Urban Audit collection (SFSO 2018).

### Data sources

The Swiss Federal Statistical Office (SFSO) provides an inventory of land statistics datasets (SFSO 2017), namely a set of land use/land cover maps for the national extent of Switzerland, which comprise 72 base categories. Four datasets have been released for 1979/85, 1992/97, 2004/09 and 2013/18^1^, at a spatial resolution of one hectare per pixel. The pixel classification is based on computer-aided interpretation of satellite imagery, which includes special treatment and field verification of pixels where the category attribution is not clear.

The SFSO land statistics datasets have been used to produce a series of categorical maps for each urban agglomeration and time period. In order to process the SFSO datasets in an automated and reproducible manner, an open source reusable toolbox to manage, transform and export categorical raster maps has been developed in Python (Bosch 2019b). The boundaries of each urban agglomeration have been adopted from the definitions provided also by the SFSO (2014), which comprise multiple municipalities and have been established in consideration of population density, proximity between centers, economic activities and commuting behavior. As stated above, Geneva and Basel are excluded from this study because a significant portion of their urban agglomeration lies beyond the extent covered by the SFSO land statistics inventory, namely the administrative boundaries of Switzerland. The spatiotemporal evolution of the urban footprint for the three selected urban agglomerations (i.e., Bern, Lausanne and Zurich) over the study period (i.e., 1980-20161) is displayed in Figure 2.

**Fig. 2.**
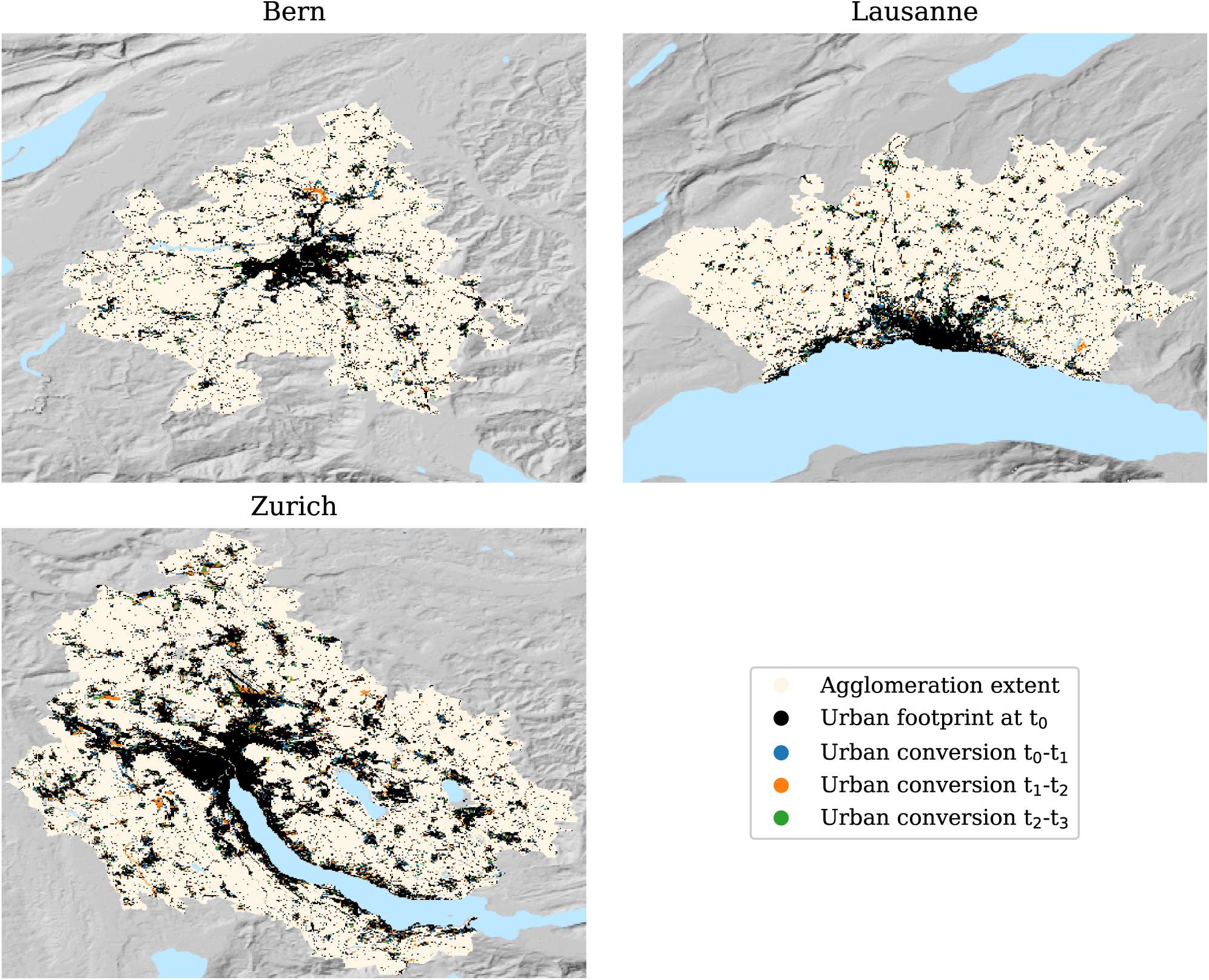
Evolution of urban patches of the three urban agglomerations throughout their respective periods of study. The times t_0_, t_1_, t_2_ and t_3_ correspond to 1981, 1993, 2004 and 2013 for Bern; 1980, 1990, 2005 and 2014 for Lausanne and 1982, 1994, 2007 and 2016 for Zurich

### Quantifying spatiotemporal patterns of urbanization

While a plentiful collection of landscape metrics can be find in the literature, many of them are highly correlated with one another. As a matter of fact, Riitters et al. (1995) found that the characteristics discerned by 55 prevalent landscape metrics could be reduced to only 6 independent factors. On the other hand, landscape metrics can be very sensitive to the resolution and the extent of the maps. However, several metrics empirically exhibit consistent responses to changing scales that conform to predictable scaling relations (Wu et al. 2002; Wu 2004). Based on such remarks, and in order to enhance comparability with other studies, ten landscape metrics have been selected for the present study, whose details are listed in Table 1. While complying with the FRAGSTATS v4 definitions (McGarigal et al. 2012), the landscape metrics have been computed with the open source library PyLandStats (Bosch 2019a). Like in most of the related studies, the categorical maps have been reclassified into urban and natural classes, and the metrics have computed at the urban class level, namely aggregating their values across all the urban patches of the landscape. The only exceptions are the contagion (CONTAG) and Shannon’s diversity index (SHDI), which can only be computed at the landscape level, namely considering all the landscape classes (urban and natural in the this study). Pixels that correspond to land unavailable for development, such as water bodies, have been excluded from the computation of the metrics.

**Table 1.** Selected landscape metrics to quantify the spatiotemporal patterns of urbanization. A more thorough description can be found in the documentation of the software FRAGSTATS v4 (McGarigal et al. 2012)

| Metric name (abbrev.) | Level | Category | Description |
| --- | --- | --- | --- |
| Percentage of landscape (PLAND) | Class | Area and edge | Percentage of landscape, in terms of area, occupied by patches of a given class |
| Mean patch size (MPS) | Class and landscape | Area and edge | Mean patch size |
| Largest patch index (LPI) | Class and landscape | Area and edge | Percentage of the landscape occupied by the largest patch |
| Patch density (PD) | Class and landscape | Aggregation | The number of patches per area unit |
| Edge density (ED) | Class and landscape | Area and edge | Sum of the lengths of all edge segments per area unit |
| Area-weighted mean fractal dimension (AWMFD) | Class and landscape | Shape | Mean patch fractal dimension weighted by relative patch area |
| Mean euclidean nearest neighbor distance (ENN) | Class and landscape | Aggregation | Mean patch shortest edge-to-edge distance to the nearest neighboring patch of the same or different class |
| Landscape shape index (LSI) | Class and landscape | Aggregation | Standardized ratio of edge length to area |
| Contagion (CONTAG) | Landscape | Aggregation | Metric that measures the extent to which patches of the same class are spatially aggregated in the landscape |
| Shannon's diversity index (SHDI) | Landscape | Diversity | Metric that measures the diversity of patch types determined by both the number of different patch types and the proportional distribution of area among patch classes. |

### Modes of urban growth

In addition to the conventional landscape metrics, which are computed over a single snapshot of a landscape, Liu et al. (2010) proposed a quantitative method to classify the types of urban growth occurring between two time points. To that end, for each new urban patch, the Landscape Expansion Index (LEI) is computed as^2^:

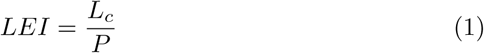

where *L*_*c*_ denotes the length of the interface between the new urban patch and pre-existing urban patches, and *P* is the perimeter of the new urban patch.

Then, the type urban growth attributed to a new urban patch will be identified as infilling when *LEI >* 0.5, edge-expansion when 0 *< LEI* ≤ 0.5 and leapfrog when *LEI* = 0. As suggested by Li et al. (2013), the relative dominance of each growth mode between two landscape snapshots can be evaluated both in terms of number and the area of new urban patches that are attributed to each growth mode.

### Fractal aspects of urban patterns

Despite their complex and irregular appearance, cities comply with well-defined order principles that can be characterized quantitatively by means of fractal geometry (Frankhauser 1994; Batty and Longley 1994). Two main characteristics of fractal structures are of particular interest in the context of the study of urbanization and land use change (White et al. 2015): the area-radius scaling and the size-frequency distribution of urban patches.

### Area-radius scaling

The relationship between the built-up area of an urban agglomeration and the distance from the main city center has been shown (e.g., Frankhauser (1994)) to empirically conform to relationships of the form:

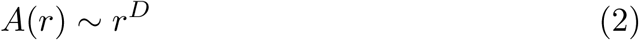

where *A* denotes the total area of the urban built-up extent that lays within a distance *r* from the city center, and *D* corresponds to the radial dimension, analogous to the fractal dimension of two-dimensional complex objects such as Sierpinski carpets. Although the measure might not be appropriate for urban agglomerations with multiple important centers, Frankhauser (1994) found extensive evidence that most contemporary cities could be approximated through (2), with the value of *D* consistently falling between 1.9 and 2.0. On the other hand, following initial observations by Frankhauser (1990), White and Engelen (1993) suggested that the area-radius scaling could be better approximated throguh two values of *D*, a first steeper one for small values of *r*, reflecting an inner zone where urbanization was essentially complete, and a second lower slope for the outer zone that is still undergoing urbanization.

### Size-frequency distribution of urban patches

Contemporary urban agglomerations are configured by numerous patches of urban land use. If such configuration is fractal, there must be no characteristic patch size, and thus the relationship between the size of an urban patch and the number of patches of that size found in the agglomeration must follow a power-law scaling relationship of the form:

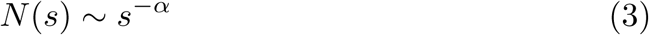

where *N* (*s*) is the number of patches of size *s*, and the scaling exponent *α* is the patch size-frequency dimension (White and Engelen 1993).

## Results

### Time series of landscape metrics

The computed time series of landscape metrics for the agglomerations of Bern, Lausanne and Zurich are displayed in Figure 3 (see also Code S1). As suggested by previous research (Wu et al. 2002; Wu 2004), the chosen landscape metrics show predictable responses to changes to the map extent (see Code S2). Therefore, the reminder will only consider the values computed at the extent of the urban agglomeration.

**Fig. 3.**
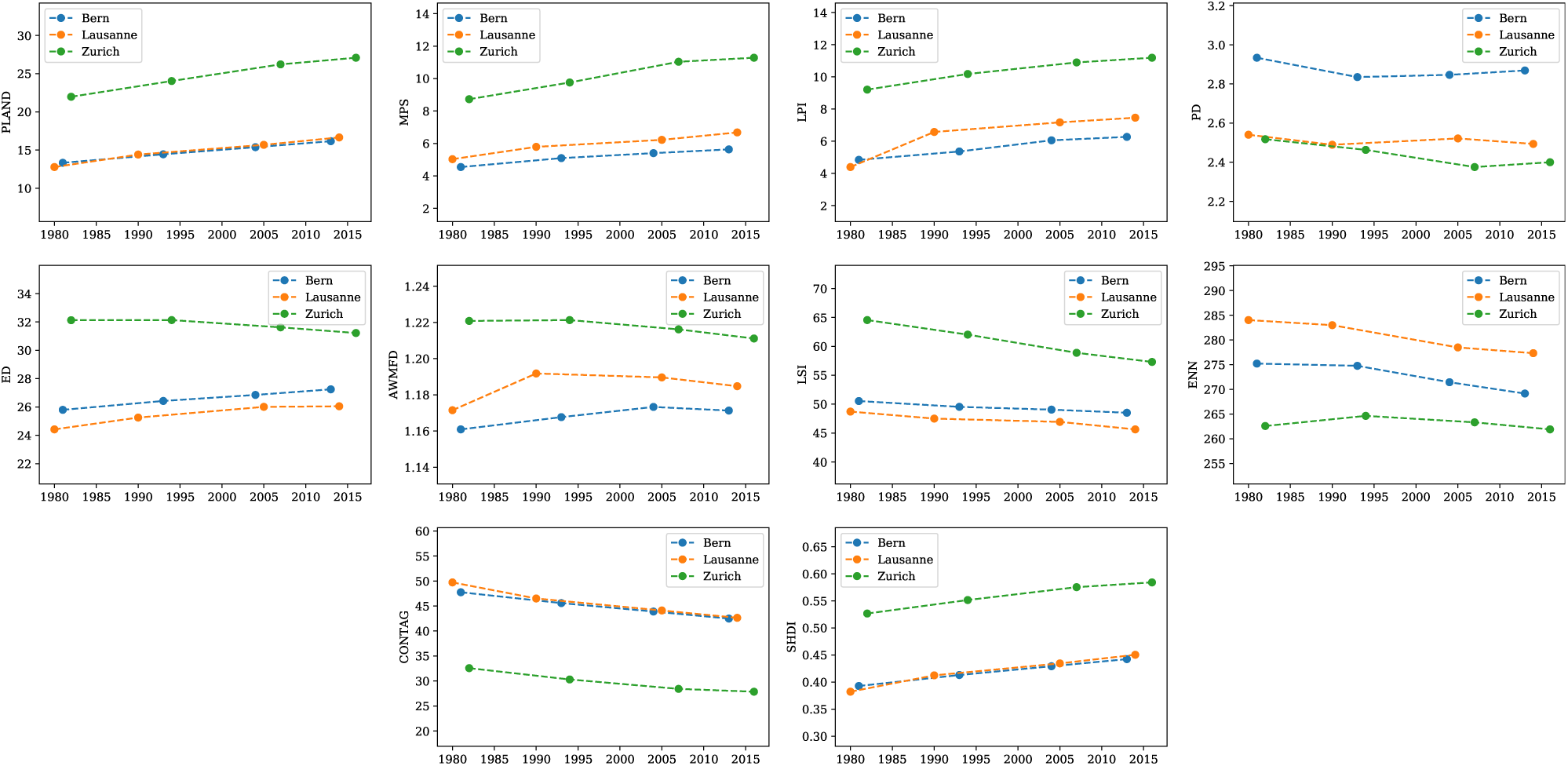
Time series of landscape metrics. The eight metrics of the two upper rows are computed at the urban class level, that is, aggregating the values computed for each urban patch. The two metrics of the lower row are computed at the landscape level, that is, considering the patches of all the classes present within the landscape (urban and natural in this case)

The proportion of landscape occupied by urban patches, represented by PLAND has increased monotonically for the three agglomerations. Bern and Lausanne show almost indistinguishable trends, starting from a 13% in the early 1980s and surpass the 16% in the last snapshot of 2013 and 2014 respectively. Zurich shows a parallel tendence with the percentage of urbanized land increasing from 22% in 1982 to a 27% in 2016. Similarly, MPS and LPI also show a monotonic increase, parallel among the three agglomerations, but again with higher values for Zurich. Such trends are overall characteristic of coalescence stages and the higher values observed in Zurich are consistent with its higher population density.

On the other hand, PD reveals more complex and idiosyncratic trends, which suggest that new urban patches can basically emerge at any period. The Bern agglomeration shows the highest PD values, which is consistent with its MPS and LPI being the lowest among the three agglomerations and denotes a landscape configured by numerous small urban patches. Despite the irregularity of PD, ED shows more consistent tendencies of increase in Bern and Lausanne and decrease in Zurich. From the perspective of the diffusion and coalescence hypothesis, Bern and Lausanne are seemingly undergoing a diffusion stage, whereas the decrease of Zurich is more characteristic of coalescence. The flat evolution of ED during the first period in Zurich suggests a transition from diffusion to coalescence, which is also observed in Lausanne during the last period. This postulate is also supported by the trend of AWMFD, which in Zurich is almost identical to that of ED, and shows an unimodal pattern for Lausanne, also characteristic of a transition from diffusion to coalescence. The decline of the AWMFD observed during the last period in Bern also suggest that albeit tardier than its counterparts, Bern might also be undergoing a shift towards coalescence. The monotonic decrease of the LSI reveals that the landscape is becoming less edgy, which is also characteristic of coalescence. Similarly, ENN shows an overall monotonic decrease which suggests that distances between neighboring urban patches are decreasing.

The two metrics that operate at the landscape level, CONTAG and SHDI show consistent monotonic trends which are almost identical for Bern and Lausanne. The decrease of CONTAG indicates that urban and natural patches are becoming increasingly disaggregated and interspersed, which is characteristic of diffusion and contrasts with the hallmarks of coalescence exhibited by most of the other metrics. On the other hand, the increase of SHDI denotes an increasing compositional diversity. The fact that the values of CONTAG and SHDI are respectively lower and higher in Zurich is mostly due to its higher proportion of urbanized landscape, which make the proportional abundance of urban and natural pixels more even than in Bern or Lausanne. Overall, given that only two classes (i.e., urban and natural) are considered within this study, CONTAG and SHDI are almost perfectly correlated with PLAND (see Code S1), because the relative abundance of each class is the main determinant of the aggregation, interspersion and diversity of its patches (McGarigal et al. 2012).

### Growth modes

The relative dominance of infilling, edge expansion and leapfrog development during the period of study is displayed in Figure 4. As suggested by Li et al. (2013), all three urban growth modes act simultaneously. Edge-expansion is the most dominant growth mode and maintains its importance throughout all agglomerations and time periods, nevertheless, it tends to diminish over time. Such a decrease of edge-expansion is compensated by an increasing influence of infilling. Leapfrog is by far the least dominant growth mode, and its influence does not show any clear trend of increase or decrease.

**Fig. 4.**
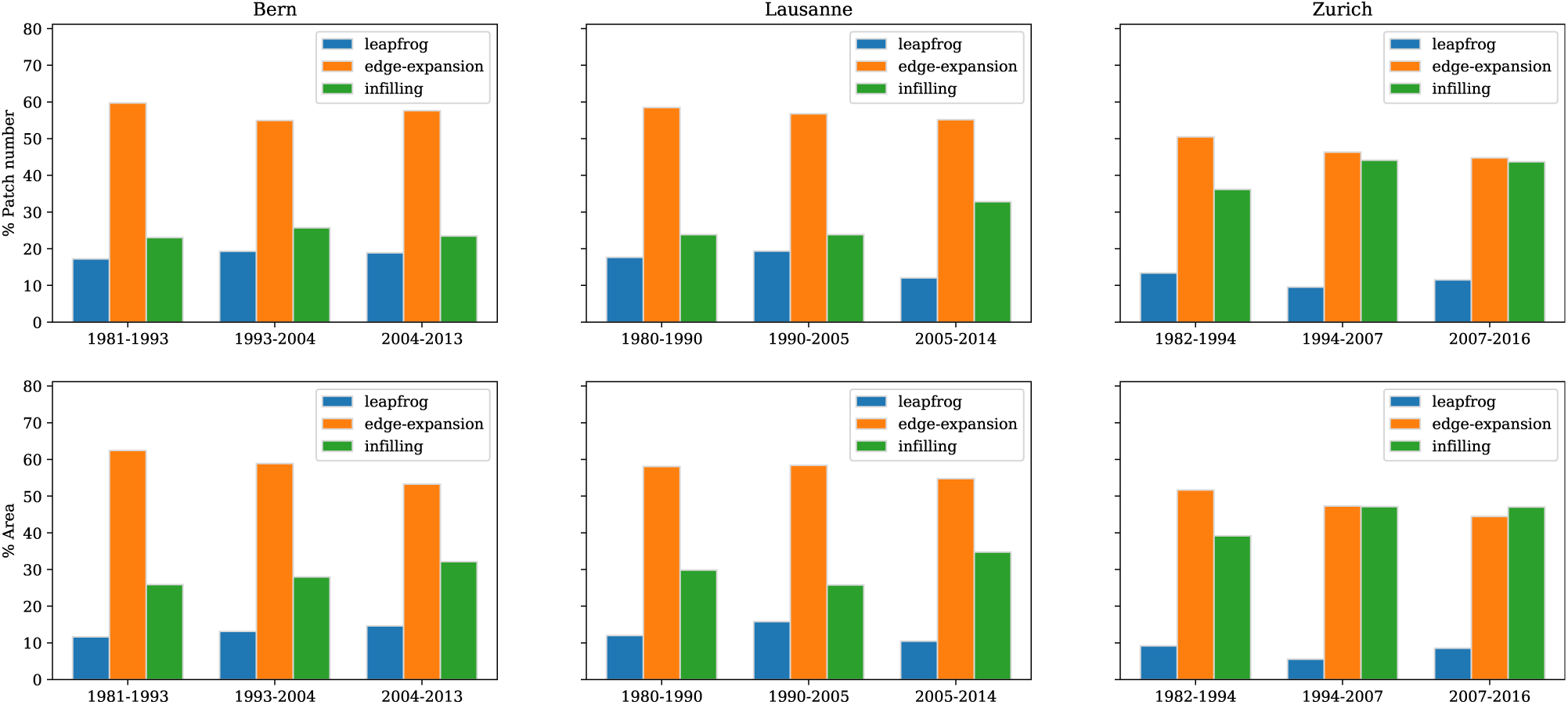
Changes in the relative dominance of inflling, edge expansion and leapfrog over the three time periods of each urbanagglomeration in terms of number of new urban patches (upper row) and their respective area (lower row).

As with landscape metrics, Bern and Lausanne show similar characteristics, namely an unequivocal dominance of edge-expansion and a slight increase of the area-weighted influence of infilling. On the other hand, leapfrog has very little influence in the agglomeration of Zurich, while the prevalence of edge-expansion and infilling is equally significant, especially in the latter periods. Like the time series of landscape metrics suggest, the higher dominance of infilling observed in Zurich is characteristic of coalescence. Similarly, the increasing influence of infilling in Bern and especially in Lausanne are consistent with the transition from diffusion to coalescence that the concave trends of ED and AWMFD seemingly indicate.

### Fractal aspects of urban patterns

The area-radius scaling and the patch size-frequency distribution of the three urban agglomerations at each temporal snapshot are plotted in Figure 5 and Figure 6 respectively.

**Fig. 5.**
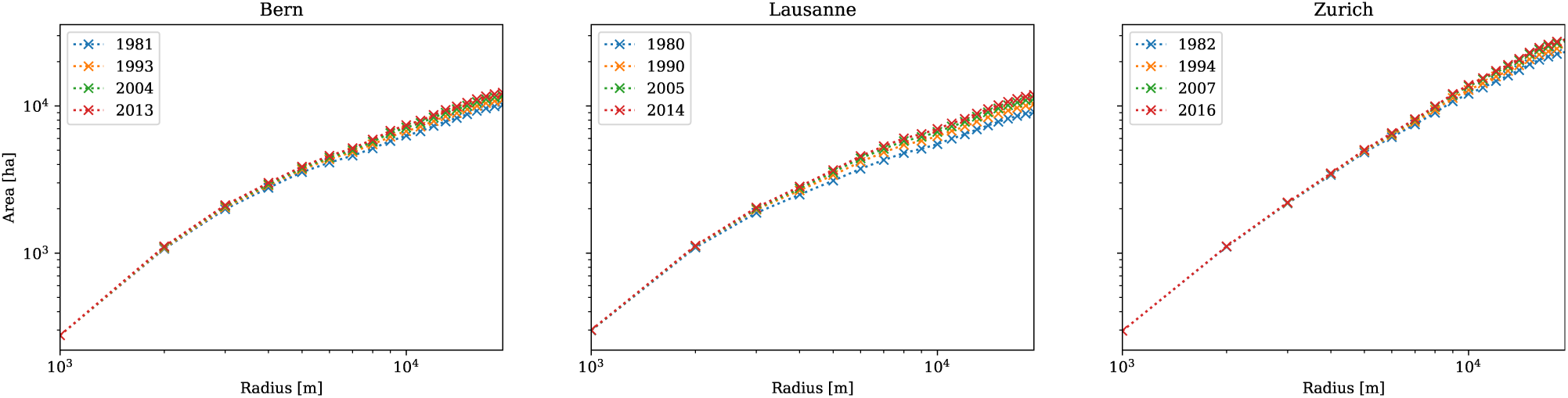
Area-radius scaling of the three urban agglomerations at each temporal snapshot. The relationship has been estimated by computing the total area occupied by urban land uses laying within a series of radius values (noted by the cross-shaped markers) from 1000 to 20000m, successively increasing by a step of 1000m. The reference center point has been manually retrieved from the OpenStreetMap^4^. See Code S3.

**Fig. 6.**
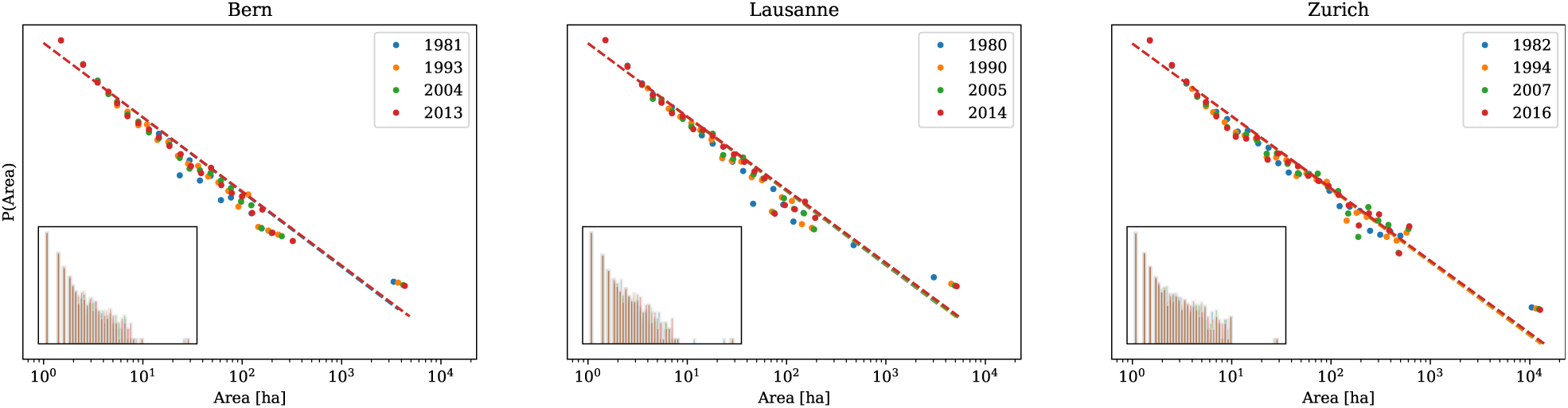
Size-frequency distribution of urban patches for the three urban agglomerations at each temporal snapshot. Each dot represents an observed patch, with the color code denoting the temporal snapshot to which the patch corresponds. The colored dashed lines represent the best maximum likelihood fit to a power-law distribution for all the patch sizes observed at their corresponding temporal snapshot. The insets display an histogram with logarithmic bins of the urban patch size distribution. The plots have been produced with the Python package powerlaw (Alstott et al. 2014). See Code S4.

In line with the observed landscape metrics and growth modes, the urban agglomerations of Bern and Lausanne show similar scaling behavior, with a kink (located around the 3000m radial distance) separating the steeper inner urbanized zone and the outer zone with more non-urban land. The area-radius scaling of Zurich is characterized by a steeper slope with practically no appreciable kink, which denotes higher proportion of urban land uses at greater radial distances from the city center. At the same time, the curves become steeper through time in all agglomerations, which reflects how they fill the available space as urbanization unfolds. Overall, the area-radius scaling curves of Bern and Lausanne are consistent with the bifractal radial city model (White and Engelen 1993; White et al. 2015). However, the temporal evolution of their scaling curves suggests that as they become more urbanized, the kink might attenuate, leading to the almost straight line observed in Zurich. Such an hypothesis seems further supported by the trends of the landscape metrics and growth modes reported above, which suggest that the kink in the area-radius scaling curve might be characteristic of agglomerations whose urban patches are still undergoing diffusion, whereas straight lines might correspond to more consolidated agglomerations whose urban pathces are coalescing.

On the other hand, the size frequency distribution of urban patches can be approximated by a power-law (see Code S4), i.e., a straight line in the log-log scale, with its slope reflecting the scaling exponent *α* of (3). The slopes of the fitted curves are very stable through time for the three urban agglomerations, although they display a slight tendence to decrease over time (note the smaller slope of the fit curves of latter years in Figure 6 and Code S4). Such a trend reflects how urban patches are becoming larger, which is in consonance with the increase of MPS observed for all the urban agglomerations. Nonetheless, the overall increase of MPS is counterbalanced by the emergence of new urban patches of smaller size, which seemingly contribute not only to keeping the size-frequency distribution of urban patches as a power-law, but also to scaling exponents that remain very stable through time. This is relatively surprising, especially considering that PD — the metric that reflects the number of urban patches — and the dominance of leapfrog growth show the most irregular trends.

## Discussion

### Testing hypothesis of urbanization patterns

The examination of the spatiotemporal patterns of land use change of the three Swiss urban agglomerations by means of landscape metrics, growth modes and fractal scaling reveal novel connections between the frameworks, which offer complementary perspectives. Firstly, the time series of landscape metrics generally support the diffusion and coalescence hypothesis, with Zurich, the largest Swiss urban agglomeration, exhibiting most characteristics of coalescence, whereas Bern and Lausanne are seemingly undergoing a transition between diffusion and coalescence. Additionally, the trend of SHDI, which is not explicitly considered in the diffusion and coalescence formulation of Dietzel et al. (2005), suggests that the landscape is becoming increasingly diverse in land use. However, it must be noted that SHDI is not very informative when considering only urban and natural classes, since it will be strongly correlated with the proportion of landscape occupied by urban patches (PLAND).

Secondly, the temporal changes in the relative dominance of infilling, edge-expansion and leapfrog are overall consistent with the characterization of urbanization as a “spiraling process that involves three growth modes of leapfrogging, edge-expanding and infilling [where] leapfrog and infilling tend to alternate in their relative dominance while edge-expansion is likely to remain its importance throughout much of the urbanization process” (Li et al. 2013). Such formulation does not necessarily contradict the diffusion and coalescence model of urban land use change. Instead, the diffusion and coalescence dichotomy might be viewed as a particular case where the way in which leapfrog and infilling alternate their dominance is not aleatory but rather characterized by a progressive decrease of the former and an increase of the latter. In any case, the importance of leapfrog growth in the present study does not necessarily decrease over time, instead it seems that infilling is becoming increasingly influent at the expense of edge-expansion. Lastly, the three urban agglomerations clearly conform to the reviewed fractal morphological hallmarks, namely the bifractal area-radius scaling and the power-law size-frequency distribution of urban patches. In coherence with the landscape metrics and growth modes, Bern and Lausanne show very similar traits. On the one hand, both urban agglomerations display a clear kink in the area-radius scaling, while the area-radius scaling in Zurich practically delineates a steeper straight line, suggesting that urbanization in Zurich fills the available space more intensely even at higher distances to the main center. On the other hand, the power-law decay as patch sizes increase is faster for Bern and Lausanne than for Zurich, which denotes that the latter urban agglomeration is dominated by larger urban patches.

Altogether, the only metric that does not conform to the diffusion and coalescence dichotomy is CONTAG, which should increase as patches coalesce, yet monotonic decreases are observed for the three urban agglomerations. Such behavior suggests that urban and natural patches are becoming increasingly disaggregated and interspersed, which might result from a combination of three factors. The first, which has already been noted above, is that the value of CONTAG strongly depends on the proportion of landscape occupied by each land use, especially when comprising only two classes like in the present study. Secondly, increases in MPS are proportionally less important than its corresponding decreases in ED and AWMFD, which is consistent with a central property of fractal objects, namely that their border is over-proportionally lengthened with respect to their area. Finally, the small yet persistent action of leapfrog growth and the irregular trend of PD are evidence that new urban patches do emerge at all periods. This suggests that fragmentation of natural patches occurs despite the apparent trend of coalescence exhibited by the other metrics. Furthermore, the continuous fragmentation of natural patches not only contributes to the decrease of CONTAG, but is also key to keeping the size-frequency relation of urban patches as a power-law with very stable scaling exponents. Given that many related studies report continuous fragmentation of natural land at the urban fringe, it is likely the size-frequency distribution of urban patches in their regions of study shows similar power-law scaling behaviors.

### Examining the persistence of fragmentation

As urbanization unfolds, urban patches are expected to grow, which suggests that eventually urban patches coalesce and thus the small ones disappear. Yet this contrasts with the continuous fragmentation of natural patches and the persistence of the power-law scaling of the urban patches observed in the present study. As a matter of fact, the complexity sciences offer a number of high-level models that intend to explain the emergence and persistence of fractal structures and power-law distributions, such as Zipf’s principle of minimal effort (Zipf 1949), or the concept of self-organized criticality (Bak et al. 1988). Nevertheless, convincingly linking the observed urban patterns to models that account for socioeconomic, cultural and political drivers of urbanization remains a challenge (Batty and Torrens 2001; Manson and O’Sullivan 2006).

The continuous fragmentation and persistence of the power-law scaling reported above might result from multiple factors. On the one hand, several key topological properties of urban street networks, such as the number of intersections per street or street length, also exhibit power law distributions (Jiang 2007), reflecting a scale-free hierarchical organization with a few major arterial roads and a large number of small streets. Since the organization of these networks exerts a central influence in the flows of people, goods and information that underpin the spatial development of cities and regions (Bettencourt et al. 2007; Batty 2008), the fact that the size of urban patches also follows a power law certainly appears as more than a mere coincidence. Furthermore, the spatial evolution of urban street networks have also been characterized by two elementary processes, namely exploration, which expands the network towards previously non-urbanized areas, and densification, where new segments bridge gaps between existing streets (Strano et al. 2012). The way in which exploration is observed to be more active in earlier stages and is followed by a phase of densification is seemingly analogous to the diffusion and coalescence hypothesis as well as to the increasing dominance of infilling. Overall, the foregoing findings urge deeper exploration of the connection between the evolution of transportation networks and land use patterns.

Another factor that calls for consideration is the decentralized organization of the Swiss spatial planning system. The urban agglomerations examined in this study are configured by multiple municipalities which might adopt distinct approaches to spatial planning. During the last decades, most of the large Swiss municipalities have adopted land management measures that incentivize the densification (Rudolf et al. 2018), yet the above evidence shows that the majority of population growth and land conversion has occured in small and mid-sized municipalities that are located further away from the agglomeration core. It is therefore very likely that such municipalities have designated new building zones, especially for residential purposes, which contribute to the leapfrogging of urban patches surrounded by natural land, and ensure a continuous feed of small urban patches that keep their size-frequency distribution stable. The lack of coordination between of planning authorities is widely regarded as one of the main causes of urban sprawl (Carruthers and Ulfarsson 2002; Mann 2009), and the revision of the Swiss Federal Act on regional planning of 2013 intends to overcome such shortcoming and define a national growth management strategy based on infill redevelopment and densification.

### Planning implications

The fragmentation of natural habitats, as observed in the three Swiss urban agglomerations of this study, has been extensively linked to negative impacts on biodiversity and key ecosystem functions (Haddad et al. 2015). Nevertheless, growth management policies that misappreciate the residential preferences for low-density environments might inadvertently encourage households to relocate further away from the agglomeration centers, resulting in longer commute times and an overall increase of sprawl at the regional scale (Schwanen et al. 2004; Robinson et al. 2005). On the other hand, the interspersion of patches of natural land within urban agglomerations provide valuable ecosystem services to its residents, such as the reduction of air pollution, alleviation of maximum temperatures, absorption of storm water, noise reduction, carbon sequestration, improvement of aesthetic and cultural values as well as the preservation of ecological habitats and biodiversity (Bolund and Hunhammar 1999; Gómez-Baggethun and Barton 2013). In Switzerland, the lowest supply of such ecosystem services is found in the cantons with highest population density, hence policies encouraging densification should consider the role of natural patches to ensure a sustainable supply of ecosystem services (Jaligot et al. 2019). To that end, recent planning approaches based on fractal geometry (Yamu and Frankhauser 2015) might be exploited in order to satisfy the residential demand for green environments while explicitly considering planning objectives such as protecting natural habitats and improving accessibility to urban and natural amenities (Bosch et al. 2019).

## Conclusion

The present study combines three different approaches to study the spatiotemporal patterns of land use change associated to urbanization of three of the main Swiss urban agglomerations over four periods of study from 1980 to 2016. The results are quite consistent with the diffusion and coalescence model of urbanization, and suggest that Zurich is already immersed in the coalescence phase, whereas Bern and Lausanne seem to be at the transition between diffusion and coalescence. However, despite the characteristics of coalescence exhibited by most metrics, continuous leapfrogging of urban patches occurs in the three agglomerations, which fragments natural land and maintains the structural complexity of the landscapes over the four periods of study. Overall, the analysis of this paper shows how landscape metrics, urban growth modes and fractal scaling can be combined to obtain insights into the spatiotemporal patterns of urbanization that would be hard to obtain with any of the approaches individually. The way in which complex fractal structures are sustained despite important changes in the landscape is reminiscent of the dynamic behaviors encountered in a wide range of complex self-organizing systems, and in the context of this study, is probably related to the distinct planning policies adopted by the various municipalities that configure each urban agglomeration, as well as to residential preferences for low density environments. With the current Federal planning precepts aiming at the concentration growth within the existing urban agglomerations, local planning authorities should devote special attention to the valuable ecosystem services that the persisting patches of natural land can provide to urban dwellers.

## Supplementary Material

### Code S1

Computation of the time series of landscape metrics and exploration of their correlations over all the urban agglomerations and the whole period of study, as Jupyter Notebook (IPYNB). https://github.com/martibosch/swiss-urbanization/tree/biorxiv/notebooks/metrics_time_series.ipynb

### Code S2

Exploration of the sensitivity of the time series of landscape metrics to the spatial extent of the urban agglomerations, as Jupyter Notebook (IPYNB). https://github.com/martibosch/swiss-urbanization/tree/biorxiv/notebooks/sensitivity_extent.ipynb

### Code S3

Exploration of the area-radius scaling of each urban agglomerations over the whole period of study, as Jupyter Notebook (IPYNB). https://github.com/martibosch/swiss-urbanization/tree/biorxiv/notebooks/area_radius_scaling.ipynb

### Code S4

Exploration of the size-frequency distribution of urban patches of each urban agglomeration over the whole period of study, as Jupyter Notebook (IPYNB). https://github.com/martibosch/swiss-urbanization/tree/biorxiv/notebooks/size_frequency_distribution.ipynb

## Acknowledgments

This research has been supported by the École Polytechnique F édérale de Lausanne (EPFL).

## Footnotes

1 The exact dates of each surveying period 1979/85, 1992/97, 2004/09 and 2013/18 are determined according to the production process of the national maps and vary accross the Swiss territory (SFSO 2017)

2 The LEI definition of (1) is taken from Nong et al. (2018) and is equivalent to the initial formula proposed by Liu et al. (2010)

4 https://www.openstreetmap.org/

## Notes

#### Summary of Updates

Changed template, updated references to recently published articles

https://github.com/martibosch/swiss-urbanization

